# Identifying discriminative EEG features of Unsuccessful and Successful stopping during the Stop Signal Task

**DOI:** 10.1101/2025.04.15.648695

**Authors:** Knut A. Skaug, Esten H. Leonardsen, René J. Huster

## Abstract

The stop-signal task is often used to study inhibitory control. When combined with electrophysiological recordings, the N2 and P3 event-related potentials (ERPs) are regularly observed. Numerous studies link both amplitude and latency differences of the N2 and P3 to failed versus successful stopping. A slower N2-P3 complex when stopping fails has repeatedly been reported across many studies and found to correlate moderately with behavioral stopping speed. However, most studies rely on averaging across trials, thereby limiting the examination of trial-by-trial dynamics. In the present study, we employed different machine learning-approaches to classify successful from failed stop trials based on time-frequency single-trial EEG data. We also tested whether attenuating the slowing effect would alter classification performance. To preserve interpretability, we first identified five group-level EEG components time-locked to stopping and then used the time-frequency representation as features in different models. Our findings suggest that regularized logistic regression can reliably classify successful from failed stopping with an AUC = 0.72. Correcting for ERP latency differences did not markedly reduce overall classification (i.e., AUC = 0.71), but the model had to compensate by leveraging subtler, broadly distributed time-frequency features. Our feature importance measure indicated that a component closely resembling the N2-P3 complex contributed largely to the classification performance, producing a sparse model. Once the slowing effect was attenuated in the data, the model still retained predictive performance but had to rely on 15 times as many time-frequency features across the five components. Thus, it is likely that multiple overlapping processes unfold during stopping that influence response inhibition in addition to the N2-P3 complex. While the N2-P3 complex is consistently evoked during stopping and carry large discriminative ability, considering additional auxiliary processes might further our understanding into mechanisms underlying response inhibition.

## Introduction

The ability to flexibly navigate a changing environment requires us to control and adapt our behavior. Underlying this ability is a dynamic interplay of higher-order cognitive control functions. As these functions are pivotal in guiding behavior, understanding how such processes unfold may offer critical insight into both healthy and pathological cognition. A key cognitive control function is response inhibition, i.e., the ability to *cancel actions*, which enables us to quickly adapt our behavior to demands or goals within an ever-changing environment. A widely popular paradigm for studying response inhibition is the Stop Signal Task (SST). The SST is designed specifically to study our ability to withhold or suppress an already initiated response (Verbruggen & Logan, 2008). It is an extension of the simpler choice-reaction time task in which participants are instructed to respond as fast as possible each time a go-stimulus is shown. The key difference is that the SST presents a stop-stimulus shortly after the go-stimulus in a minority of trials. The stop stimulus instructs the participant to withhold or suppress their already initiated action towards the first-presented go-stimulus. By varying the duration between the go-and subsequent stop-stimulus, the task of stopping is made harder or easier, which in turn allows experimental control of accuracy across the stop trials. This dynamic tracking aims to produce a balanced number of successful and failed stops during a session.

The SST has been used extensively to study the underlying neural processes involved during inhibitory processing in healthy populations (Huster et al., 2013; Verbruggen & Logan, 2008), and how these processes might be disrupted in patient populations (Lipszyc & Schachar, 2010). The importance of understanding neural processes underlying response inhibition is critical as disrupted inhibitory processing may have profound effects on everyday life by causing impulsive behavior, which in turn can be detrimental for the individual (Bari & Robbins, 2013). Understanding the dynamics and mechanisms underlying response inhibition may also help dissociate how different clinical symptoms might be driven by disrupted inhibitory processes.

Much research has therefore been devoted to identifying signatures of response inhibition to build an understanding of how these relate to specific aspects of stopping an action.

A longstanding influential formal model describing the processes involved in going and stopping in the SST is the horse-race model (Logan & Cowan, 1984). Simply put, the process of going and stopping is modelled as a “horse-race” between two independent processes. The process that wins the race determines whether the outcome is successful or unsuccessful. Because a successful outcome is marked by the absence of a response (no button press), it cannot be measured directly, and a model of the processes unfolding is needed to estimate the time of the unobserved response. The horse-race model accomplishes this by estimating how long it takes on average to cancel the response towards the go-stimulus, i.e., the stop-signal reaction time (SSRT). This metric is often used as a behavioral index of inhibitory control, and individual-or group-level differences in response inhibition are often equated with differences in the SSRT. However, recent concerns question the reliability of the SSRT as a stable marker of individual differences in inhibitory control (Thunberg et al., 2024). Others have demonstrated that group differences reflected in the SSRT might be inflated by some individuals experiencing an objectively harder task and that controlling for objective task difficulty reduces observed brain activation differences (D’Alberto et al., 2018). While the SSRT has been used extensively to contrast patients from control populations (Verbruggen & Logan, 2008) its reliability may be limited or confounded by objective task difficulty. Therefore, identifying stable markers that enable a more reliable characterization of inhibitory processing differences is crucial. This need motivates the search for reliable markers beyond a behavioral composite score like the SSRT.

Because the race unfolds rapidly across milliseconds, electroencephalography (EEG) has been used extensively to study the temporal dynamics of response inhibition (Huster et al., 2013, 2020; Wessel & Aron, 2015). EEG markers of response inhibition occur after the stop stimulus and typically comprise the stereotypical N2 and P3 components at fronto-central electrodes (Huster et al., 2020). Researchers have long debated whether the N2 and P3 truly mark processes involved in cancelling the motor command to the go stimulus (Hervault & Wessel, 2024; Wessel & Aron, 2015), or if they rather reflect the detection/evaluation of an unexpected or conflicting event (Huster et al., 2020; Skippen et al., 2020). However, they usually agree that there is a timing difference between failed and successful stopping, with failed stopping lagging successful (Huster et al., 2020; Hervault & Wessel, 2024), an empirical finding which is also predicted by the race-horse model (Logan & Cowan, 1984). The P3 peaks around 300ms while the N2 peaks around 200ms, and some believe the P3 is associated with slower (2-3Hz) oscillations whereas the N2 is slightly faster and associated with 4-5Hz oscillations (Huster et al., 2017; Huster et al., 2020). Common criteria used to identify potential EEG markers of response inhibition seem to be that their EEG signatures differ between (1) *stopping and going* (Huster et al., 2017), (2) *successful and failed stopping* (Bekker et al., 2005; Kok et al., 2004; Ramautar et al., 2004; Schmajuk et al., 2006; Skippen et al., 2020; Wessel & Aron, 2015), and ideally that (3) *successful versus failed stopping differences should covary with the standard behavioral index defined as the SSRT* (but also see Huster et al., 2020, for a more extensive list of criteria).

In the case of the N2 and P3, several studies report larger amplitudes during stopping than going (Huster et al., 2013; Kok et al., 2004). When comparing successful and failed stopping, results are more mixed. Some report larger N2 during failed stopping (e.g., (Ramautar et al., 2004; Wüllhorst et al., 2025), while others find no difference (Kok et al., 2004) or suggest N2 amplitude reflects infrequent or salient events more generally (Enriquez-Geppert et al., 2010).

Numerous studies report larger P3 amplitude for successful stopping (Bekker et al., 2005; Greenhouse & Wessel, 2013; Wüllhorst et al., 2025). A delay in either N2 or P3 latencies during failed stopping is more consistent across studies (Bekker et al., 2005; Ramautar et al., 2004; Wüllhorst et al., 2025). Lastly, both the N2 and P3 latencies seem to reliably correlate with the SSRT (Huster et al., 2020).

Some argue that contrasting successful vs. failed stopping is overly conservative as it is likely that the same brain networks are active during both outcomes just to a different degree (Boehler et al., 2010). However, separating unique processes driving successful or failed stopping is necessarily best achieved through comparing successful and failed stop trials. Thus, when performance is defined behaviorally as successful vs. failed stopping, EEG-based markers sensitive to inhibitory processing should consistently characterize this contrast across individuals.

Besides the N2-P3 complex, multiple studies suggest that another electrophysiological signature of stopping is increased beta power over right frontal scalp areas (Hannah et al., 2020; Wagner et al., 2018; Wessel, 2020). Transient beta bursts increase during successful stopping (Wagner et al., 2018; Wessel, 2020) but also to unexpected events (Tatz et al., 2023; Wagner et al., 2018), suggesting a possible role that likely involves sensory processing of infrequently presented stimuli, like the stop signal. Furthermore, unique trial-by-trial patterns of bursts predict reaction times (Little et al., 2019; Wessel, 2020), a relationship likely obscured using traditional averaging techniques given its transient dynamics. Importantly, both the N2-P3 complex and the beta-burst features of stopping have been found in studies employing spatial filtering to produce source-level scalp projections (Hervault et al., 2024; Wagner et al., 2018), demonstrating that spatial decomposition is able to disentangle reliable features evoked during stopping. Spatial filtering methods have also been used to discover novel features underlying successful stopping relative to going (Huster et al., 2017), providing a new set of features enabling high discriminative ability at single-trial level. Huster et al., (2017) further demonstrated how combining single-trial features with machine learning allowed for out-of-sample predictions often unavailable in traditional ERP analyses. These powerful methods successfully identified novel features with high predictive ability, thus moving towards a more dynamic view of brain responses and how these interact to shape behavior on a trial-by-trial level. So far, to our best knowledge, few studies have applied machine learning to classify single-trial data acquired during cognitive control paradigms (Gholamipourbarogh et al., 2023; Huster et al., 2017; Vahid et al., 2018, 2020). Vahid and colleagues (2018) used a support-vector machine approach to predict behavioral performance in a Go/No-Go task. They identified novel EEG features associated with the motor cortex beyond the N2 and P3 to predict behavioral performance, but did not directly classify single-trial activity. In a later study, Vahid et al., 2020 employed deep learning to classify single-trial EEG data from a Simon task, achieving classification accuracy of approximately 33% above chance level for discriminating conflict from non-conflict trials. Gholamipourbarogh and colleagues (2023) used EEG data from a Go/No-Go task, combining group-level spatial decomposition with deep learning, and identified four distinct spatial topographies present when individuals had to inhibit their responses. Importantly, each contributed significantly to the model’s predictive ability. Moreover, the topographies unfolded during similar time periods indicating that multiple processes likely happen simultaneously, which might not be captured by traditional univariate approaches. Collectively, these studies demonstrate that machine learning approaches allow for a rigorous data-driven discovery of neurophysiological features at the single-trial level and that they can capture simultaneous cognitive control processes that might otherwise be missed by traditional univariate approaches.

Because overt behavioral responses in the SST only occur when stopping fails, identifying reliable neurophysiological signatures is critical for uncovering the processes differentiating successful from failed stopping. Extending the scope beyond the typically estimated N2-P3 complex and beta-burst dynamics at fronto-central scalp areas may uncover new EEG markers of inhibitory processing that provide a clearer and a more complete picture of response inhibition.

Although prior studies demonstrate the usefulness of combining spatial filtering with machine learning to disentangle important features underlying inhibitory processing, no prior study has contrasted successful from failed stopping at single trial level in the SST. We therefore set out to explore whether these methods could be applicable to discriminate successful from failed stopping, possibly contributing novel signatures which would help progress the field forwards. Our goal in this study was first to identify possibly novel discriminative features underlying successful and failed stopping in the SST by applying group-level spatial decomposition methods alongside machine learning to classify successful from failed stopping at single trial level. Our second goal was to evaluate whether we could apply these methods to test already established signatures. Specifically, we wanted to test the idea that failed stopping is largely marked by slower inhibitory processing (Huster et al., 2020, Wessel & Aron, 2015). To test this, we conducted the same analysis on two versions of the same dataset and evaluated model performance and stability using cross-validation. Specifically, for one of the datasets, we minimized the inter-trial latencies across all trials prior to decomposition and modeling. As we did this across both successful and failed trials combined, it effectively attenuates temporal processing differences when individuals fail to inhibit their response.

## Methods

### Participants

A total of 45 participants took part in this study which included two sessions of EEG recording separated by two weeks. 38 participants completed both sessions, leaving a total of 82 individual datasets. We excluded 9 individual datasets because the class balance between successful and failed stopping exceeded a threshold set to 65%. A total of 40 participants were included in the final analysis of which 32 contributed two individual datasets (two sessions) while 8 participants only had a single dataset (one session). Due to an error, one dataset was included twice in the analysis. Provided that 72 of 73 datasets were unique, the possibility of substantially biased results due to this doubling is minimal.

### Experiment

The entire EEG recording session comprised the SST, a Stroop task, a Go/NoGo task, a task-switching task, and finally a resting period with eyes open after completing all tasks. The experiment was presented on an Eizo FlexScan S2411W monitor connected to a Dell Precision T5500 computer (Dell, Inc., Texas, USA), and programmed using PsychToolbox-3 (version 3.06.16) in MATLAB R2019a (The MathWorks, Inc., Massachusetts, USA). Behavioral responses were collected using SuperLab RB-740 (Cedrus Corporation, 2006). Because the focus in this study is on the SST, only this task will be described. The SST comprised a blocked design presenting pseudo-randomly drawn go-and stop-trials with a total of 450 trials. Each block comprised 68 go trials and 22 stop trials, yielding a stop signal probability of 24%. All trials started with a black fixation cross before either a single go-stimulus or a successive pair of go-and stop-stimuli were presented. All stimuli were centered in the middle of the screen. The fixation cross was presented for a duration of 700-1200ms whereas the go-or stop-stimuli had a duration of 100ms. These go-and stop-stimuli comprised orange or blue arrows pointing either to the left or the right, indicating which hand to respond with. During stop trials, the color of the second arrow (i.e., the stop stimulus) changed relative to the first (i.e., go stimulus) instructing the participants to refrain from responding. The order of possible color combinations was counterbalanced across participants. Stair-wise SSD tracking with a range of 100-600 ms changed dynamically in 50 ms steps. Performance feedback followed each block instructing the participants to be faster, more accurate or both.

### Data Acquisition

EEG was recorded using a BrainAmp system (Brain products GmbH) with 31 passive channels (Ag/AgCI) placed according to the international 10-20 system in an electrically shielded room. The data was sampled at 5000 Hz, referenced to channel FPz with the ground placed at AFz.

Impedance was below 5 kΩ for all channels.

## Data analysis

### Preprocessing

All preprocessing was carried out using the EEGLAB toolbox (v2022.1; Delorme & Makeig, 2004) and custom-made in-house preprocessing scripts in MATLAB 2024a (The MathWorks, Inc., Massachusetts, USA) (see appendix for more details). Continuous data were down-sampled to 512Hz, high-pass filtered above 0.75Hz and low-pass filtered below 40Hz. Independent component analysis (ICA) was carried out to project out components identified as muscular or ocular using the EEGLAB toolbox “IClabel” with thresholds set at 0.35 for both classes of artifacts. Channels containing excessive noise were identified and spherically interpolated before the data were re-referenced to the average reference. Local artifact-correction was next carried out to remove residual large and focal artifacts remaining after artifact correction by ICA. Finally, any remaining bad channels were again identified and spherically interpolated.

### Post processing

To test the hypothesis that latency differences represent an important feature underlying successful and unsuccessful stopping, a second data set was generated with attenuated latency differences (i.e., the latency-corrected dataset). This was done by correcting for latency jitter across trials including both successful and unsuccessful trials, thus attenuating latency differences between both conditions (see appendix for details). Both the main and modified datasets were then epoched to 600ms pre-stimulus and 600ms post-stimulus. All subsequent procedures were then carried out separately for the modified and main dataset (see Figure 1 for an overview). We corrected for imbalances between classes (SS or US) by randomly selecting a subset of trials from the largest class to match the number in the smallest class. We excluded individual datasets which initially had a class imbalance ratio exceeding 65:35 from further analysis as this was too extreme to be corrected. The same individual datasets were excluded for both the modified and the main dataset. Next, we estimated spatial filters across both trial types using group task-related component analysis (*for specific details into this method see*: (Tanaka, 2020)). We added a Thikanov regularization to the covariance matrices to ensure they were full rank and better conditioned. This was done in Matlab as such: 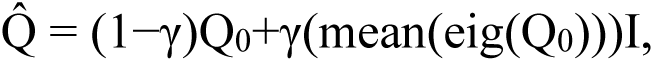 where γ was set to 0.1, Q_0_ is the covariance matrix, and I is the identity matrix. The top five eigenvectors were extracted for each individual dataset and normalized to unit vectors. Forming spatial filters across both conditions should ensure that no bias was introduced across experimental conditions (Cohen, 2022). Spatial maps were calculated by mean-centering and normalizing each eigenvector to unit variance, then multiplying the data with the normalized component and dividing the resulting product by the number of time points. Due to the inherent sign uncertainty of eigenvectors, the sign of the electrode with the largest absolute value was used to either multiply with-1 if the sign was negative or retain as it was if the sign was positive (Cohen, 2022). To compute the temporal components, we normalized the eigenvectors to unit length and multiplied with the data matrix. If the spatial map was multiplied with-1 the temporal component was similarly multiplied with-1. This procedure reduced the number of channels to five components for each individual dataset yielding a component x time x trial matrix. Finally, each component’s time x trial matrix was transformed into a time-frequency representation using Morlet wavelet decomposition (Cohen, 2019). Specifically, flipped copies of the first and last trial were appended to the epoched matrix, and each trial was zero-padded with half the epoch length attached to each side to attenuate edge-effects. Wavelets were constructed for 60 logarithmically spaced frequencies between 2-30Hz, with the number of cycles increasing on a logarithmic scale from 2 cycles at the 2Hz to 12 cycles at 30Hz. The trial matrix was vectorized before frequency-domain convolution. After transformation, the extra padding and the wavelet’s half-length was trimmed and the vector was reshaped back into its original epoched shape. The absolute magnitude was calculated for each single trial yielding a component x frequency x time x trial matrix. Importantly, time-frequency amplitudes are insensitive to signal polarity arising from sign-uncertainty making comparison across datasets valid.

**Figure 1.**
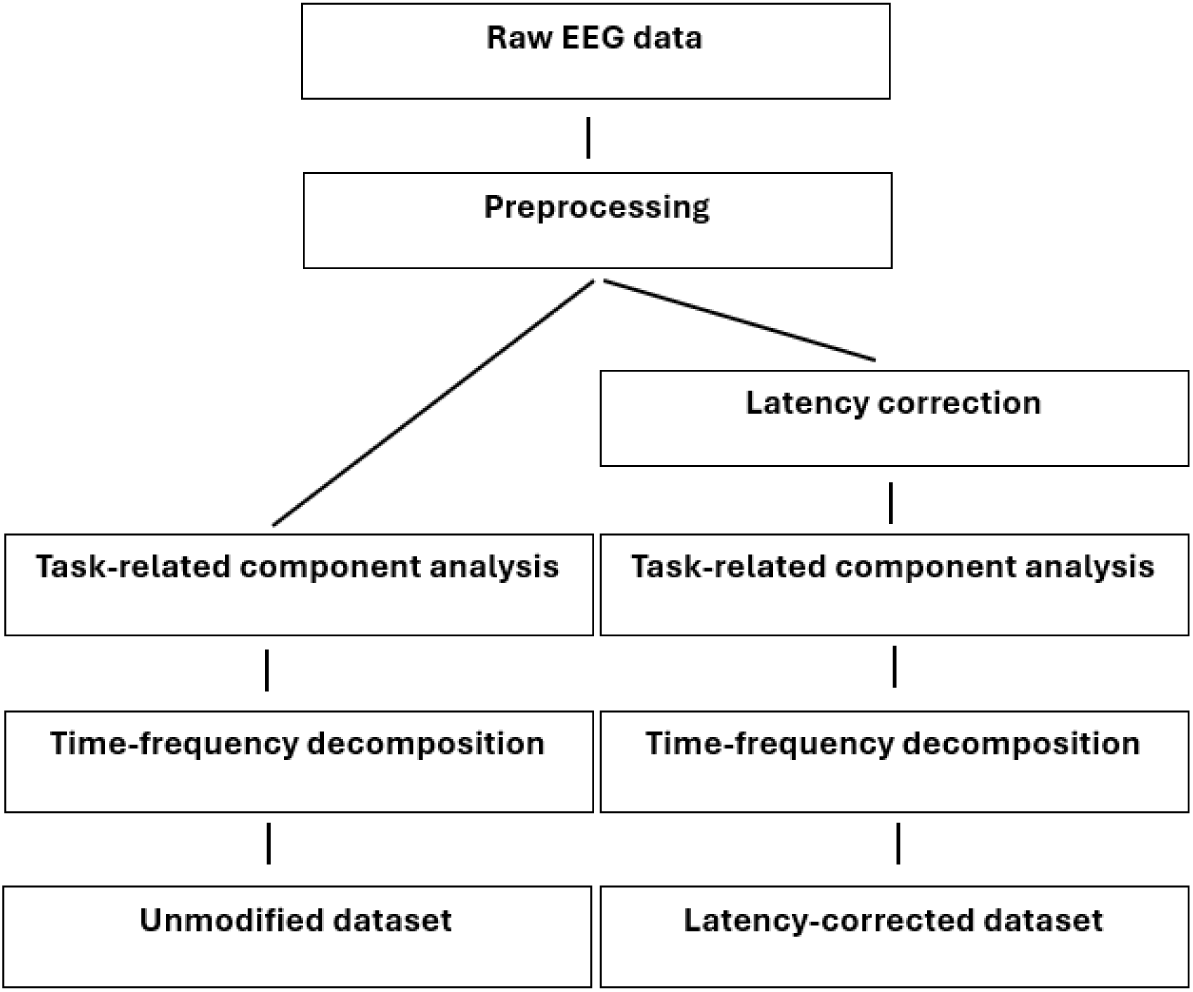
Signal processing steps

### Statistical modelling

We fit predictive models to differentiate between the successful and unsuccessful trials based on processed time-frequency single-trial data. For both datasets independently, we fit binary classifiers of four different kinds to assess what type of model most accurately captured the predictive signal: Standard logistic regression models, multi-layer perceptrons (MLPs), bi-directional long-short term memory networks (LSTMs), and gradient boosted trees (XGBoost). For the logistic regression, MLP and XGBoost models, we flattened the 3-dimensional input array (time x frequency x component) to form a singular input vector. For the LSTMs, we retained the time-dimension to match the time-steps of the model and flattened the remaining two dimensions to achieve a vector at each time-step.

Next, for each model class, we fit multiple models to span a range of reasonable hyperparameters. For each model configuration, e.g. concrete combination of model class and hyperparameters, we fit five models in a cross-validation loop, for each iteration using different portions of the dataset to train and test the models (Figure S1). Specifically, we split the full dataset into five disjoint subsets, referred to as folds. These folds were constructed on the level of sessions, such that for each iteration of the cross-validation, all trials from a single session resided either exclusively in the training or the validation set. To determine the best model, we computed the mean Area under the Receiver Operating Characteristic curve (AUC) across the held-out validation folds.

For the logistic regression models, we employed an *l*_1_-loss and varied the regularization parameter (λ ∈ {1*e*^−3^, 1*e*^−2^,…, 1*e*^3^}) and the number of max iterations (*n* ∈ {100, 300}). For the MLPs we varied the weight decay (η ∈ {0, 1*e*^−3^}), the dropout rate (*d* ∈ {0, 0.3}), the number of hidden layers (*l* ∈ {3, 5}), and the number of artificial neurons in the first hidden layer (*v* ∈ {64, 256}). The subsequent layers contained half the number of neurons of its predecessor. For the LSTMs we varied the weight decay (η ∈ {0, 1*e*^−3^}), the dropout rate (*d* ∈ {0, 0.3}), the number of LSTM-cells (*l* ∈ {3, 5}) and the number of neurons per cell (*v* ∈ {64, 256}). For the XGBoost models we varied the number of trees (*t* ∈ {25, 100}), the max depth of each tree (*m* ∈ {3, 7}) and both the lambda and alpha regularization-parameters, (both α, λ ∈ {0, 1}). Finally, to compare information content between the two datasets, a Wilcoxon signed-rank test was employed to determine if their distributions of performance were significantly different.

### Interpretation of coefficients

To better understand what regions of the EEG-data contained predictive information, we investigated the coefficients of the best logistic regression models for both the unmodified and latency-corrected dataset. Stacking coefficients across the five folds from the cross-validation yielded a 4-dimensional tensor with dimensions corresponding to the five folds, the five components, 60 frequencies, and 358 timesteps, yielding a tensor with dimensions folds x components x frequency x time. To remove coefficients that were not consistent across runs, we created a binary mask of shape components x frequency x time with a 1 in all locations where there were at least 3 non-zero coefficients along the dimension representing the folds, and where all non-zero coefficients had the same sign (i.e., either positive or negative). This mask was multiplied with the entire coefficient map to effectively remove ambiguous weights. Next, we collapsed the first dimension by calculating a mean coefficient across the folds. Finally, for each component, we plotted the remaining frequency x time matrix for visual inspection.

### Permutation testing for variable importance

To gain an understanding of which components were important to the models, we employed a permutation testing scheme where we fed the models partially scrambled data. The scramble was performed component-wise, such that for each location within that component (e.g. a specific time step and frequency), we randomly shuffled values across the trials and participants. This was performed for each fold individually, each time using the model for which the fold was unseen, yielding five AUCs per scrambled component. To assess whether scrambling one component resulted in significantly worse model performance than another we performed a one-sided Wilcoxon signed rank test between the AUCs of each component-pair.

## Results

In the unmodified dataset, the predictive performance of the models spanned AUCs from 0.5 to 0.72 (Figure 2a). When split according to model type, the distribution of AUCs were statistically indistinguishable for the logistic regression, MLP, and XGBoost models, whereas the LSTMs performed significantly worse than the others. Overall, the best model was a logistic regression-model with λ=0.1 and 100 iterations that reached a mean AUC of 0.72 and a mean accuracy of 65% across the folds.

**Figure 2.**
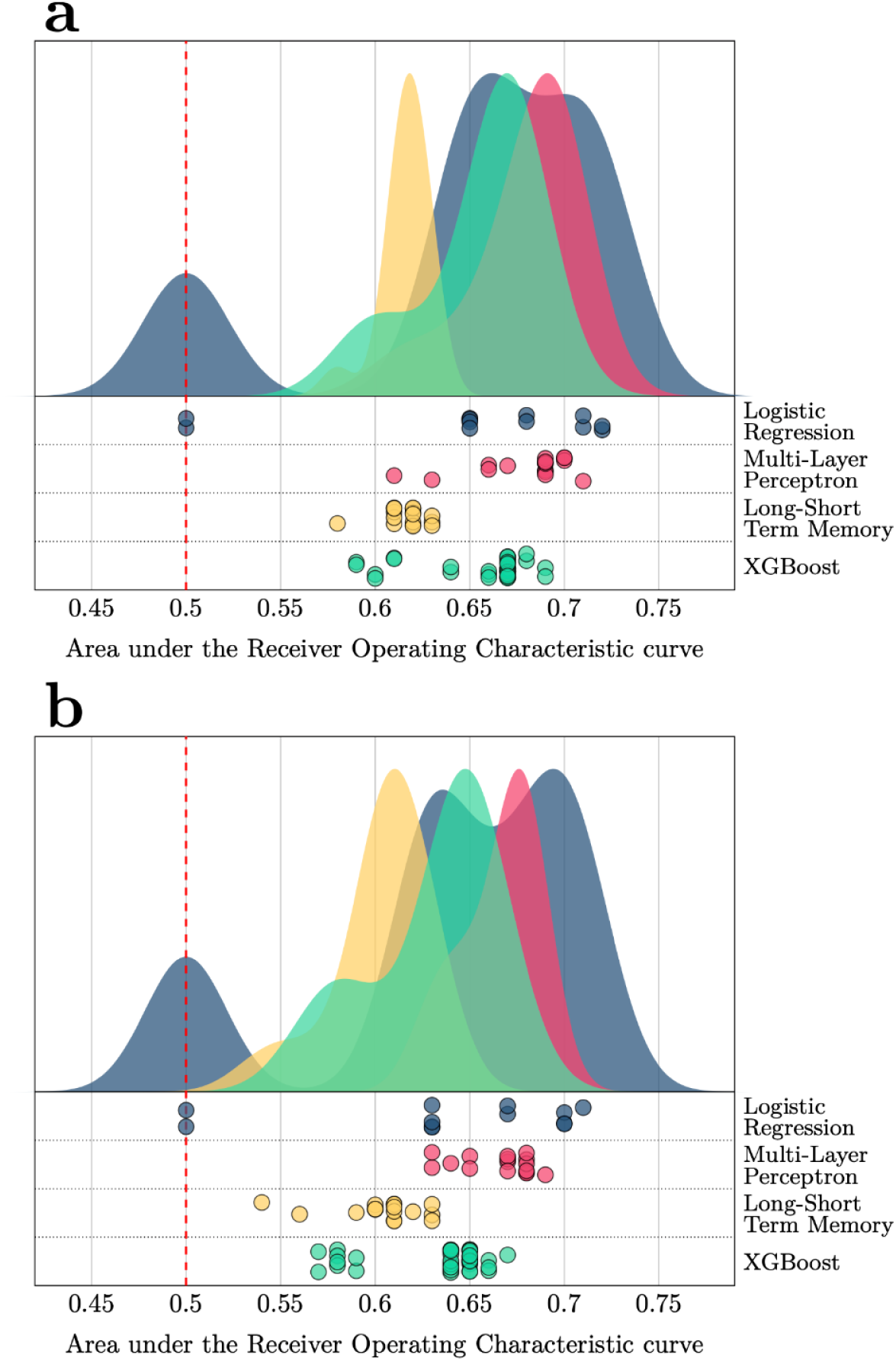
Predictive performance of the different model configurations. Note: We tested 75 model configurations, spanning four model types with various hyperparameter setups. For each configuration we performed a cross-validation, and the dots represent the mean AUC across the folds for each of these. The distributions on top are smooth density plots across these points. Note the constrained x-axis. a) AUCs using the unmodified dataset, b) AUCs using the same hyperparameter configurations in the latency-corrected dataset. The dashed red line indicates an AUC of 0.5, representing the theoretical baseline of predictive performance.

In the dataset comparison, we found that models fitted using the latency-corrected dataset resembled those fitted in the unmodified dataset, spanning AUCs from 0.5 to 0.71 (Figure 2b). The best model was also here a logistic regression-model with λ=0.1 and 100 iterations, reaching a mean AUC of 0.71 and a mean accuracy of 65% across the folds. Despite the apparent similarity between the results, when comparing results across the model configurations in a pairwise fashion, the models trained on the unmodified dataset consistently outperformed those trained on the latency-corrected data (73/76 AUCs higher or equal in the main dataset, p=4.22 × 10^−12^).

When assessing the predictive performance on the level of individual participants in the unmodified dataset, the best model achieved accuracies spreading from 52% to 82%, with a mean of 65±7% (Figure 3). 56 out of 73 participants had an accuracy better than 59.8%, the theoretical upper 95% confidence interval for a random model, and the distribution of accuracies was significantly better than random (p=4.87 × 10^−30^)

**Figure 3.**
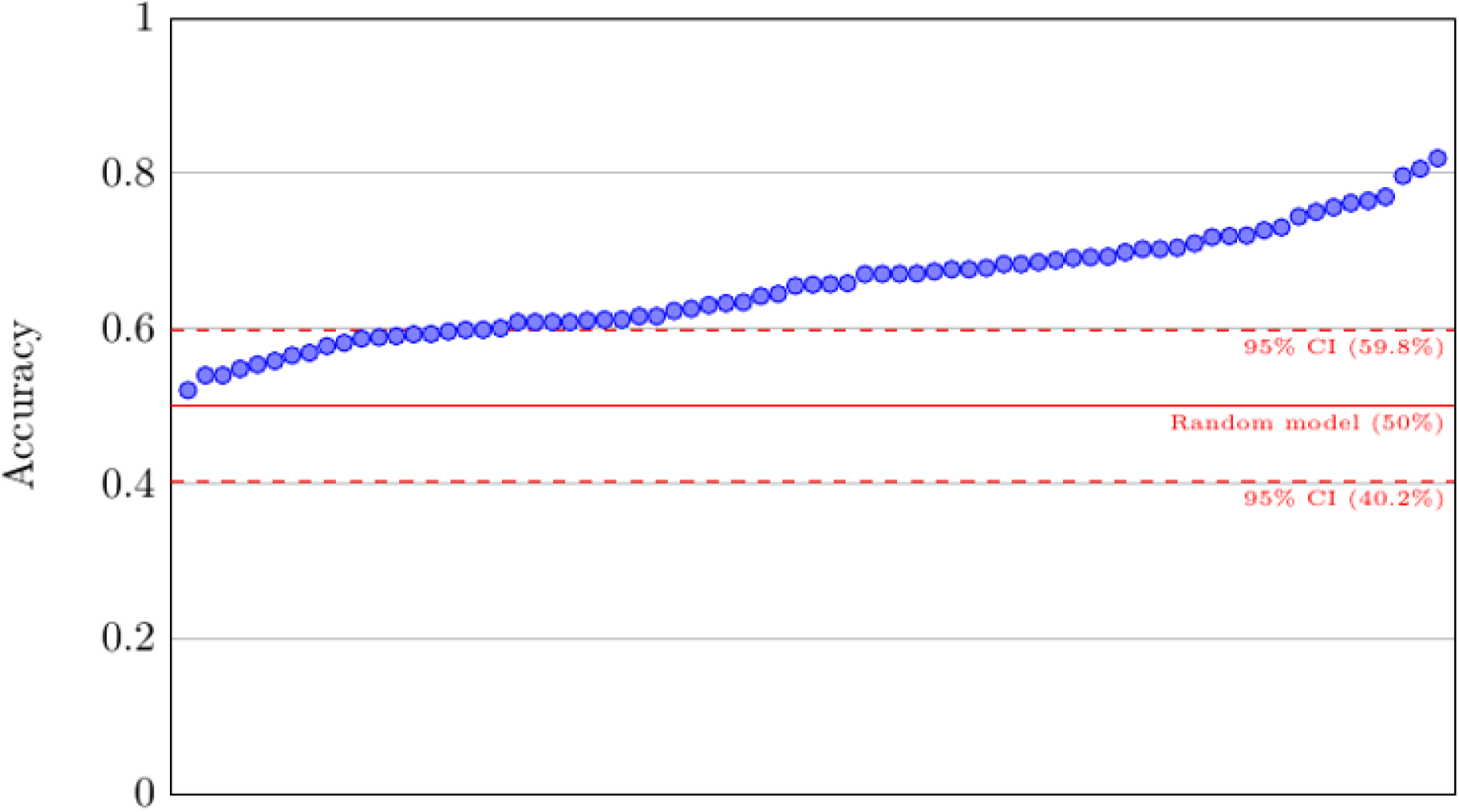
Predictive performance at the level of individual datasets. Note: Each blue dot represents the accuracy of an individual dataset across trials. The x-axis is sorted from lower to higher accuracies. For the best performing model, all datasets were above 50% chance level. The dashed red lines indicate the 95% confidence intervals for a random model.

The visualizations of the coefficients from the best logistic regression models in each of the two datasets showed clear differences, indicating that the models achieved their relatively similar performance through distinctly different predictive strategies (Figure 4d and 5d). The models trained on the unmodified dataset were very sparse, with only ∼1% (1083/107400 indices) of the coefficients passing the threshold for being considered consistent and non-zero. Most of these occurred in the first and second component (372 and 363 respectively), more than double the amount of consistent and non-zero coefficients than in the three remaining components. The coefficients with the highest absolute value, a proxy for importance, appeared in a cluster in the first component and the first frequency-band, at the very end of the time-sequence. For the latency-corrected dataset, there were substantially more consistent and non-zero coefficients (16688/107400, ∼15%), indicating that the predictive signal was spread out across the input domain to a larger degree. These were also more evenly distributed among the components, with all five components having between 2970 and 3525 consistent and non-zero coefficients.

**Figure 4.**
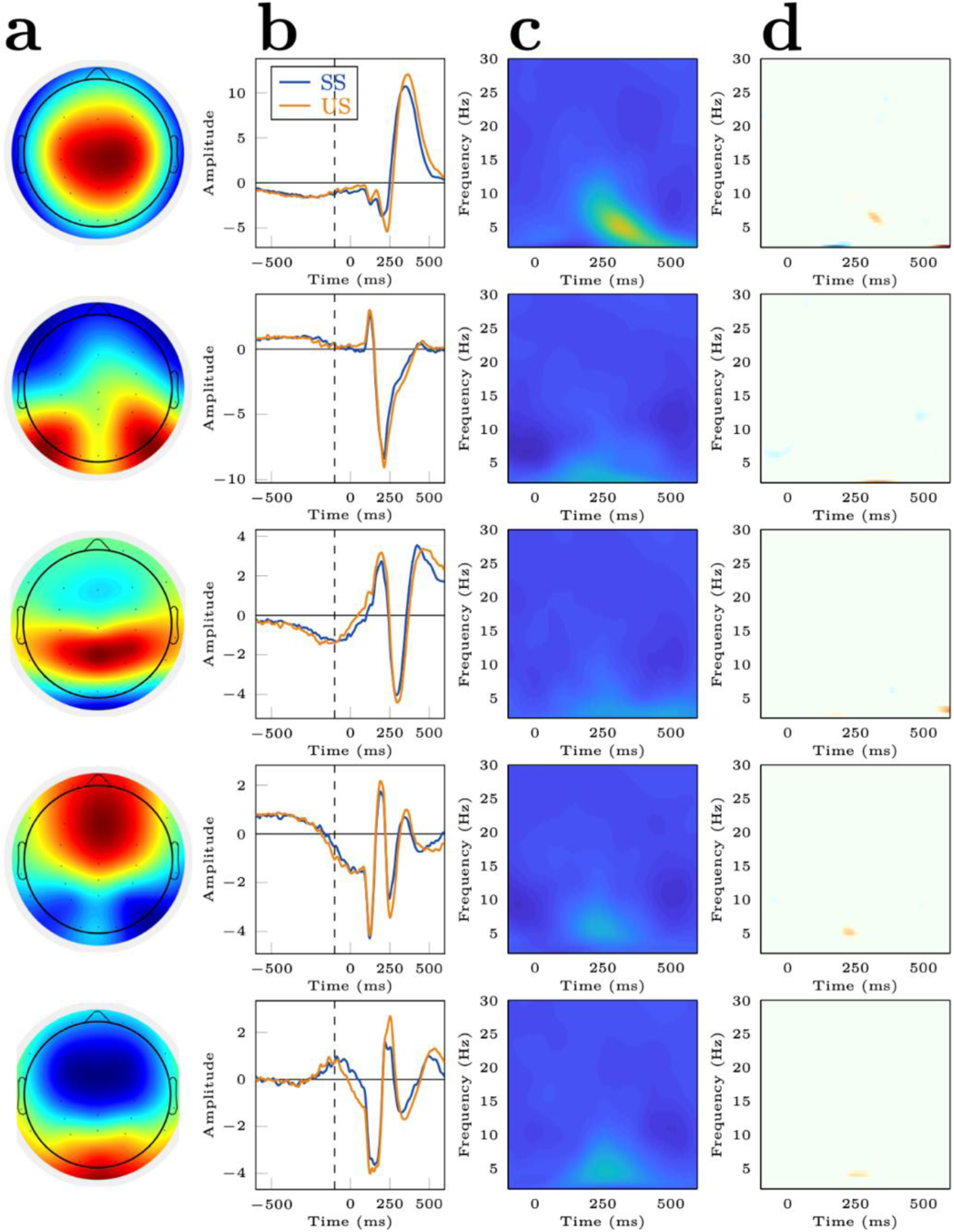
Interpretation of the differences between successful and unsuccessful trials in the unmodified dataset. Note. (a) Group-level topographies (b) Grand-average component ERPs for each condition. The dashed line indicates the start of the temporal window that was used in the modelling. To account for polarity ambiguity, the group average was flipped if its maximum was negative and individual ERPs and topographies were flipped to match the group average. This was done by comparing the summed Euclidian distances between individual (and its inverted version) and the average. Normalization to unit length ensured the validity of Euclidian distance. The smallest distance determined whether it was flipped or not. (c) Differences in the time x frequency matrix between successful and unsuccessful trials. (d) Consistent, non-zero coefficients from the logistic regression models

**Figure 5.**
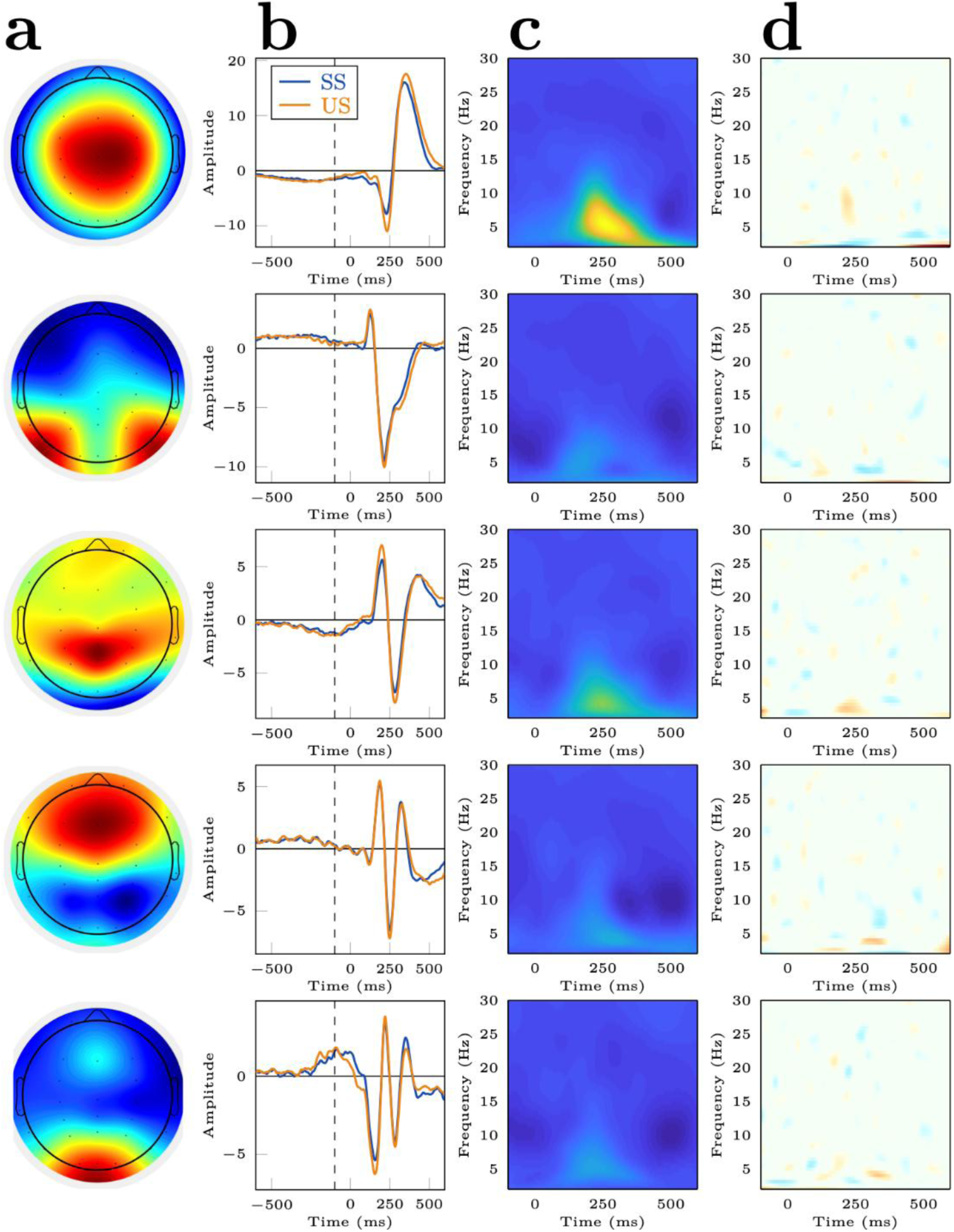
Interpretation of the differences between successful and unsuccessful trials in the latency-corrected dataset. Note. See Figure 4 for descriptions of the individual panes

However, also here the most important coefficients occurred in the first component, the first frequency-band, at the end of the time-sequence. The overall pattern emerging among the coefficients of the two datasets hinted at the unmodified dataset having a more salient and concentrated predictive signal for the models to learn, whereas in the modified dataset the models achieved the same predictive performance by finding more dispersed differences.

### Permutation importance for components

In the variable importance test we observed that scrambling any of the input components yielded a significantly worse result (p<0.05) than when using complete data (Figure 6 and Table 1), indicating that they all contained information useful to the model. The largest drop in model performance was seen when scrambling the first component (mean AUC=0.65), which was also significantly worse than the results achieved when scrambling any of the other channels, emphasizing its importance. Scrambling the second component (mean AUC=0.68) was also significantly worse than scrambling the third or fourth (mean AUCs=0.70 and 0.71 respectively), but beyond this there was no significant difference between the latter components. Overall, this corroborates the impression from the coefficient plots, i.e. that the first component was the most informative to the model, but that all channels contained information that was useful. We saw a similar picture when performing the same permutation test in the latency-corrected dataset (Figure S3 and Table S1).

**Figure 6.**
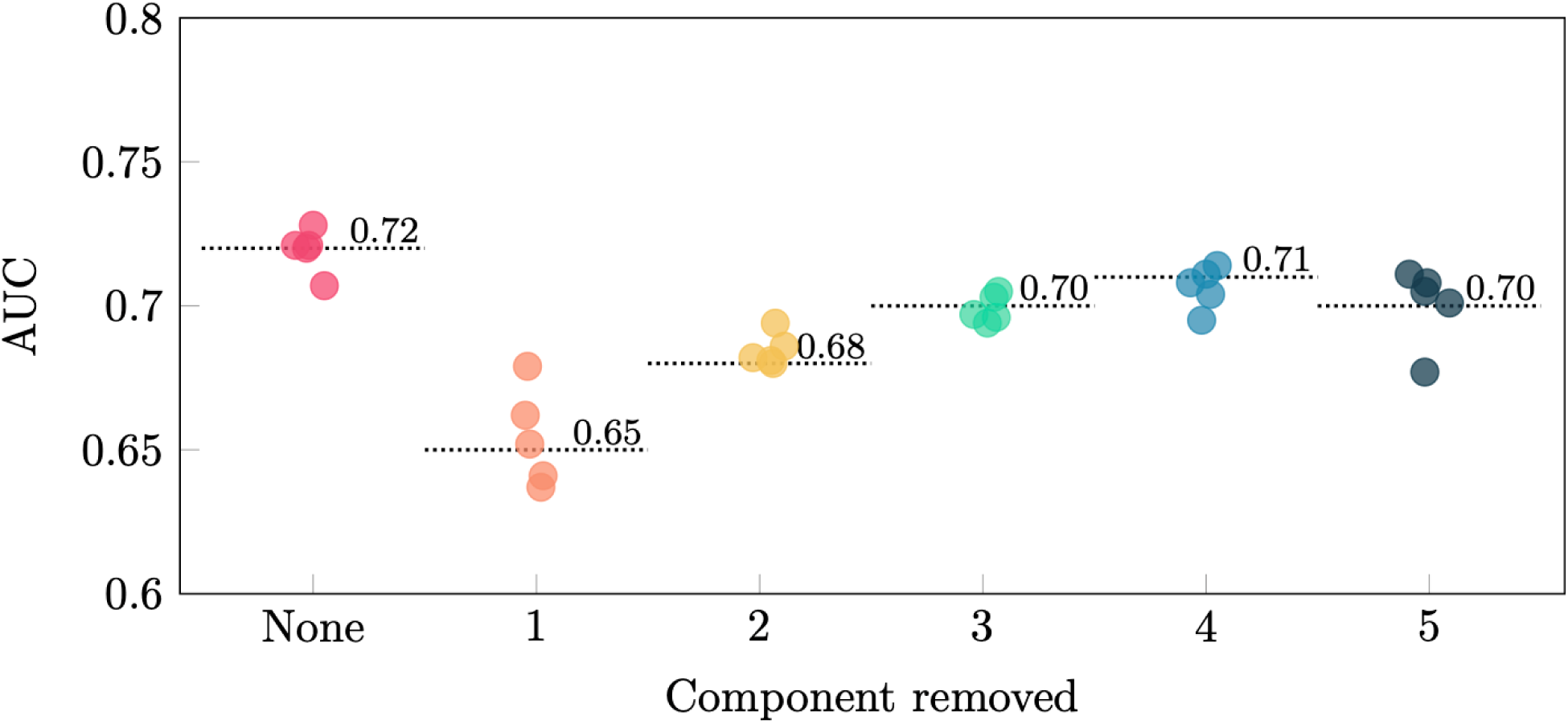
Results from the permutation test assessing the importance of individual components in the main dataset. Note: Each dot corresponds to the AUC achieved by a single model in its validation set. Each collection of dots corresponds to the performance across the five iterations of the cross-validation. The model labeled “None” refers to the original model, where no components we scrambled. The remaining five models denote performance after scrambling the contents of the given component

**Table 1.** Significance of the difference between the models removing the various components.

|  | None | 1 | 2 | 3 | 4 | 5 |
| --- | --- | --- | --- | --- | --- | --- |
| None |  |  |  |  |  |  |
| 1 | * |  | * | * | * | * |
| 2 | * |  |  | * | * |  |
| 3 | * |  |  |  |  |  |
| 4 | * |  |  |  |  |  |
| 5 | * |  |  |  |  |  |

*Note:* Each cell $[i, j]$ in the table denotes whether the models removing the component denoted by row $i$ was significantly worse than the models removing the component denoted by column $j$ , with \* indicating $p < 0.05$

## Discussion

Our results from the best performing model indicate a moderate classification accuracy across individuals for successful and unsuccessful stopping at single-trial level with an AUC around 0.72. In addition to the commonly used N2-P3 component at fronto-central electrodes, we identified four additional components, each with distinct spatio-temporal profiles, that collectively contributed to the models’ predictive power. These novel features can potentially enhance our understanding of the sources and temporal progression of EEG activity underlying successful and unsuccessful stopping in the SST. To our surprise, the simpler, interpretable logistic regression model outperformed more complex, non-linear models (bi-directional LSTMs, XGBoost, and MLPs) suggesting that linear methods capture subtle differences at single-trial level in the SST reasonably well. We also wanted to test the hypothesis that latency differences between SS and US represents an important feature for classification. Re-synchronizing the data did not influence the overall classification accuracy substantially; however, the unmodified dataset produced simpler models with fewer non-zero coefficients. This indicates that the unmodified dataset contained relationships easier to model although the overall predictive performance was similar.

Our evaluation of the importance of each component for the model’s predictive power indicated that all components contributed significantly to the overall classification performance, with decreasing importance as the component indices increase. The idea of this approach was to randomly permute one component at a time (while holding the other components constant) and evaluate the independent impact each component had on classification performance. A caveat of this approach is that generalized eigenvalue decomposition sort components by the amount of explained variability. Sorting according to variance may partly drive the observed gradient in importance, rather than reflecting intrinsic differences in the relevance of each component for classification. While taking this moderation into account, the results indicate that the first component has the largest impact on performance across folds relative to baseline and all other component permutations. The topography and temporal profile of the first component closely resemble the fronto-central N2 and P3 typically explored in electrophysiological studies of the SST. The large proportion of explained variance across trials and participants indicates that it is consistently evoked during stop signal processing. When inspecting the difference map and the grand-average ERP, frequencies corresponding to the N2 and P3 suggest both increased amplitudes and a delayed ERP in unsuccessful stop trials. Although the precise functional significance of these ERP components is still being debated, our own line of research suggests that increased N2 amplitudes might correspond to enhanced conflict detection while increased P3 amplitudes might reflect response evaluation or monitoring processes (Enriquez-Geppert et al., 2010; Huster et al., 2020). In addition to amplitude differences, a delayed response is typically observed when stopping fails (Huster et al., 2020; Wessel & Aron, 2015) and a recent meta-analysis found that this latency slowing correlates moderately with individual differences in SSRT (Huster et al., 2020). Some suggest the P3 latency might qualify as a direct marker of inhibitory processing (Hervault et al., 2024; Wessel & Aron, 2015), others suggest it is more likely an indirect marker of inhibitory processing, given its moderate correlation with SSRT and its lack of specificity (Huster et al., 2020). Our analysis set out to identify consistent features which can be exploited when contrasting successful from failed stopping on a trial-by-trial basis. Recognizing the possibility that multiple partially overlapping processes unfold whenever participants need to inhibit their responses, our results cannot confirm whether these features function as direct or indirect markers of inhibition. However, our results support the interpretation that the first component might reflect higher control engagement and evaluative processes following stopping in that the largest coefficients separating the two classes were located after N2 and P3.

After attenuating latency variability, the model still identified discriminative features. However, the model had to focus on subtler, broadly distributed EEG differences across components and time-frequency features to compensate. This illustrates a key point when interpreting logistic regression models; when a dominant discriminative feature is attenuated, the model can adapt by emphasizing other, previously overshadowed features. For the unmodified dataset, two-thirds of all non-zero coefficients were found in the first two components, and only 1% of all possible time-frequency points were utilized. For the latency-corrected dataset, approximately one-fifth of all non-zero coefficients were found in each of the five components, and it relied on 15 times as many time-frequency points as the unmodified dataset. The fact that the model retained predictive power while attenuating a dominant feature highlights the importance of other features, indicating the possibility that other processes overlapping in time can separate successful from failed stopping, although often overshadowed by the primary timing effect.

The second largest component has a topography loading maximally on posterior channels with power differences arising earlier relative to the first component. Our feature importance analysis suggests this component had only a small effect on the AUC but still contributed more to the predictive power of the models than the remaining three. It is possible this component captures early sensory processing of stimulus properties which has been shown to dissociate successful from failed stopping (Boehler et al., 2009, 2010; Knyazev et al., 2008). Boehler and colleagues (2009) additionally demonstrated that these early sensory processes are sensitive to behavioral adjustments on a trial-by-trial basis indicating perceptual/attention processes modulate performance in the SST by allocating resources towards stopping when participants successfully stop. Resource allocation through possibly proactive adjustments before the stop stimulus onset likely arise during early sensory/attentional processes (Langford et al., 2016). These results indicate that considering the impact of go-stimulus processing might provide an additional layer of information possibly influencing processes unfolding subsequently during stop stimulus processing. Although distinguishing successful from failed stopping may largely be driven by the N2-P3 component, others have found that early processing difference better predict a speed-accuracy tradeoff in the Go/No-Go task than the N2-P3 at single subject level (Stock et al., 2016). Thus, these early signals might provide an additional piece of information although their effects are likely small. It is interesting to note that non-zero coefficients weighing larger on successful relative to failed stopping are present early in our time-frequency coefficient matrix. A plausible interpretation is that there might be processes involved at or around the onset of the stop signal which carry small but discriminative information related to the behavioral outcome.

The third, fourth and fifth components all contributed minimally to classification performance as indicated by the feature importance evaluation. While the third and fifth component have a more posterior topography, the fourth component has a more frontal topography. The third component shows coefficients maximal at later time periods whereas the fourth and fifth component shows differences 200-300ms. This suggests that they possibly reflect different but overlapping processes arising during stopping in the SST. Although their contribution to classification accuracy is negligible in the main dataset, it is possible these components reflect secondary processes involved in inhibitory control. As the first component drives classification performance, their lower impact might be due to smaller effect sizes, high inter-individual variability, or temporal overlap with the first component. The time-frequency difference maps suggest most components are maximally different in the lower frequency range with largely overlapping frequency points. Interestingly, visual inspection of the third component shows that failed stopping lags successful stopping at later stages of the ERP which is also indicated in significant coefficients late in the temporal window.

Our feature importance method tests whether permuting a global predictor (i.e., component) influences the overall classification accuracy of the model. From this we cannot state whether a specific time or frequency window comprising negative or positive coefficients is more important than another area for classification. Put differently, we cannot confidently claim that the feature defined between 200-300ms in the theta range is more important for classification than another feature set. As each row in the feature matrix comprise a vectorized component x time x frequency matrix for each trial, each point is treated relative to all other, and the coefficients represent partial effects, learned in the presence of the remaining information. We cannot know how the model distributes weights if for example two features are highly correlated in two components. As such, an area with relatively small coefficient sum might still carry unique information into classification in the absence of certain other features. To test the importance of a specific feature one would need to use more local importance methods, for example, by permuting values within a feature inside one component, or permuting combinations of features. Nevertheless, the stability selection underlying our coefficient plots informs us about how consistent a specific feature appeared consistently across cross-validation folds. In other words, features with high coefficients denote a stable and strong partial effect but does not give information about whether that feature covaried with others or was unique for classification.

### Spatial-filtering and ERPs

Estimating spatial filters in task-based EEG data relies on consistently evoked activity patterns and spatial topographies across trials. Although our best performing models had moderate predictive power, it remains an open question whether reducing spatial dimensions is the best approach for identifying features in time-locked ERPs. That is, methods for blind source separation do not guarantee that projections are maximally informative for the classification problem at hand. It is possible that more complex modelling techniques are better suited for this classification problem without spatial filtering as a prior step to reduce spatial dimensions. The inherent tradeoff is that models with higher complexity often require exponentially more data and offer little interpretability into the feature-importance. Thus, we opted for a more engineered approach, acknowledging that we might not achieve the best accuracy but still retain a fairly good level of interpretability. Our focus was not to find the best classification model, but rather to identify features consistently evoked when stopping is signaled in the SST. Inspection of the average topographies suggests our components captured broadly distributed sources across midline, and lateral posterior channels.

## Conclusion

Our findings show a moderate classification of successful and failed stopping at single-trial level using EEG with a logistic regression model achieving an AUC of ∼0.72. The strongest contributor was a component resembling the N2/P3 complex, which was consistently evoked and carried high discriminative value. Attenuating latency differences between successful and failed stopping did not substantially reduce classification performance, but the model had to rely on a broader set of time-frequency features across components. This suggests that timing differences are not the sole drivers of classification. Other potentially overlapping components contributed unique, but smaller information – particularly one with posterior topography and early power differences, possibly reflecting early sensory or perceptual processes. The fact that logistic regression outperformed more complex models suggests that the identified features are linearly separable and allow for interpretation. Our approach focused on identifying repeatable EEG features related to stopping success and failure in the SST rather than aiming for maximal accuracy. Our work moves beyond trial-averaged ERPs, supporting the need for interpretability and offering a framework to study repeating processes at single-trial level which may inform future research interested in disentangling response inhibition in cognitive control.

