## Supplementary information for "Identifying discriminative EEG features of Unsuccessful and Successful stopping during the Stop Signal Task"

**Supplementary material**

**Figure S1**

*Overview of the cross-validation scheme*

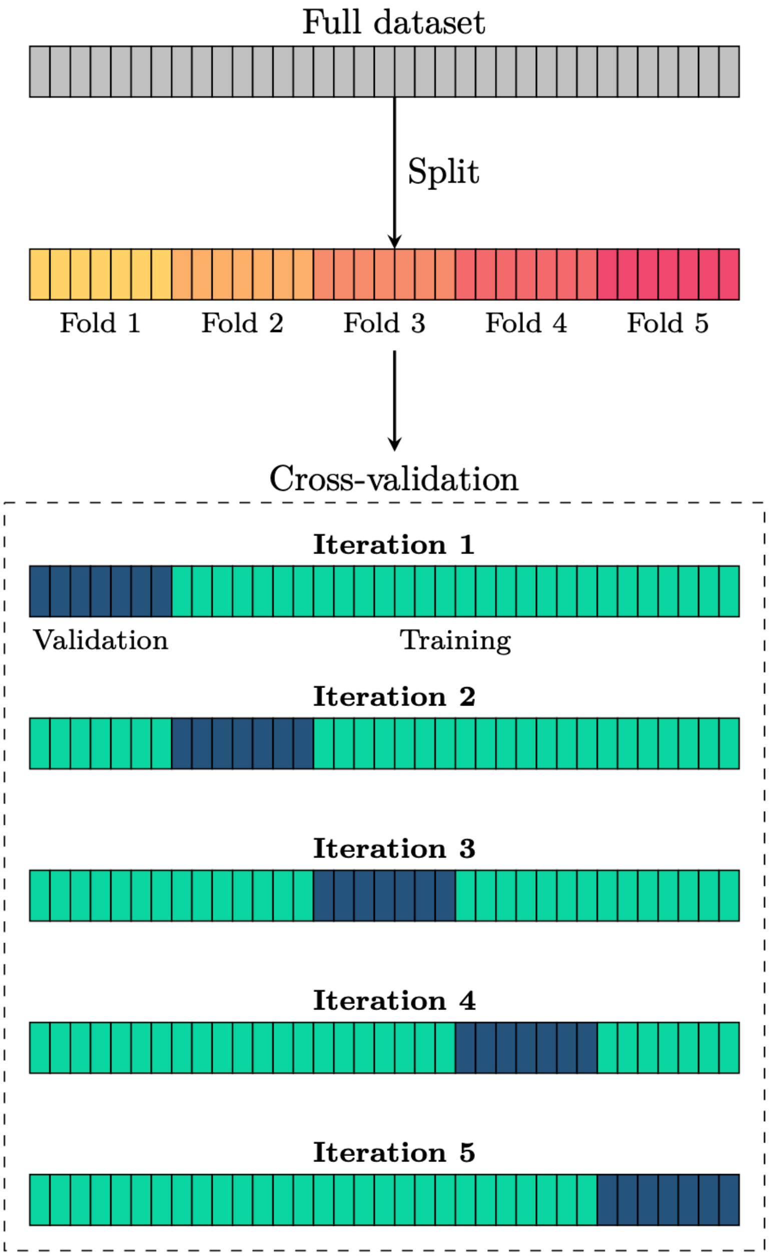

**Figure S2**

*Kernel densities of single-trial probabilities*

Latency-corrected dataset

Unmodified dataset

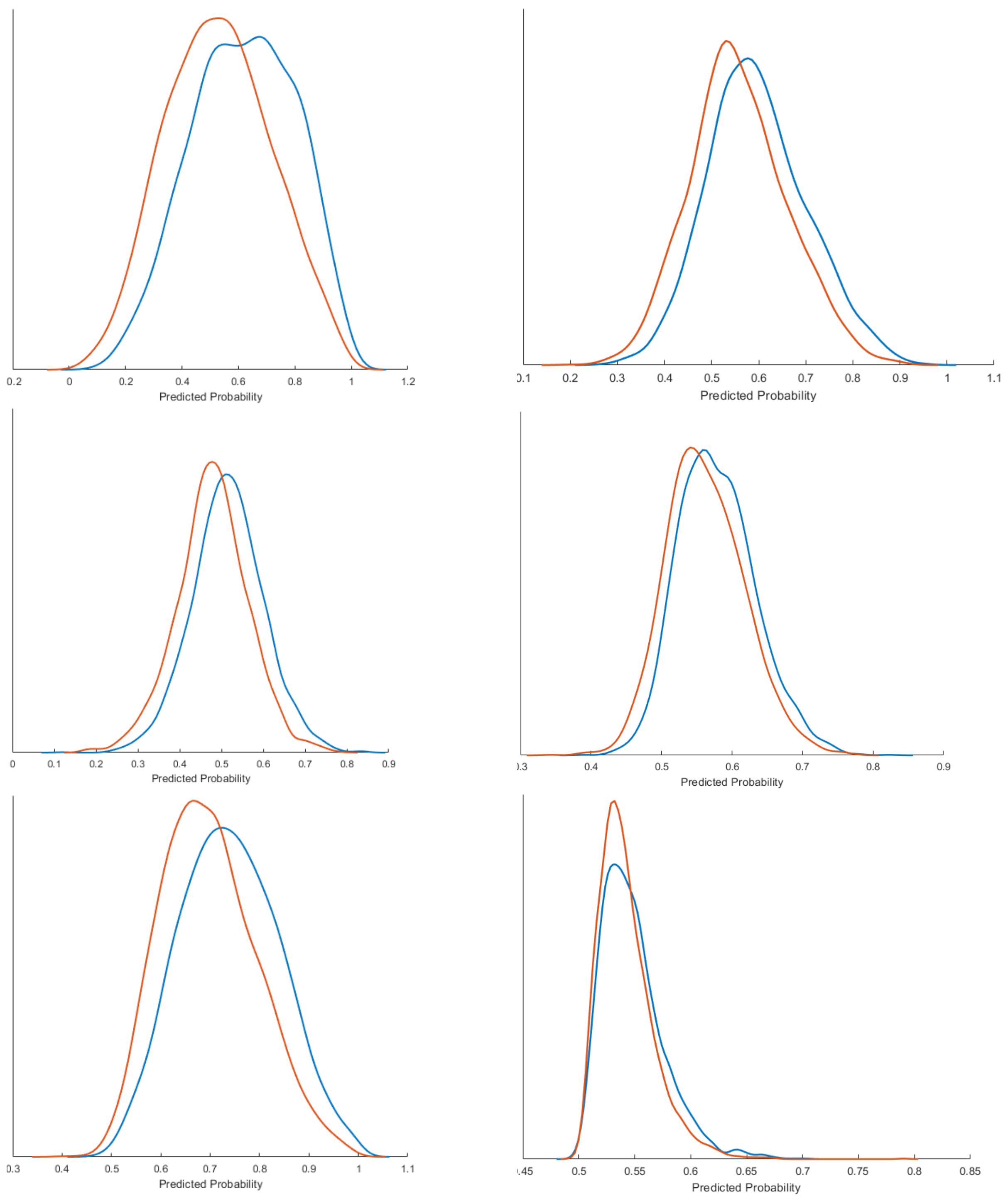

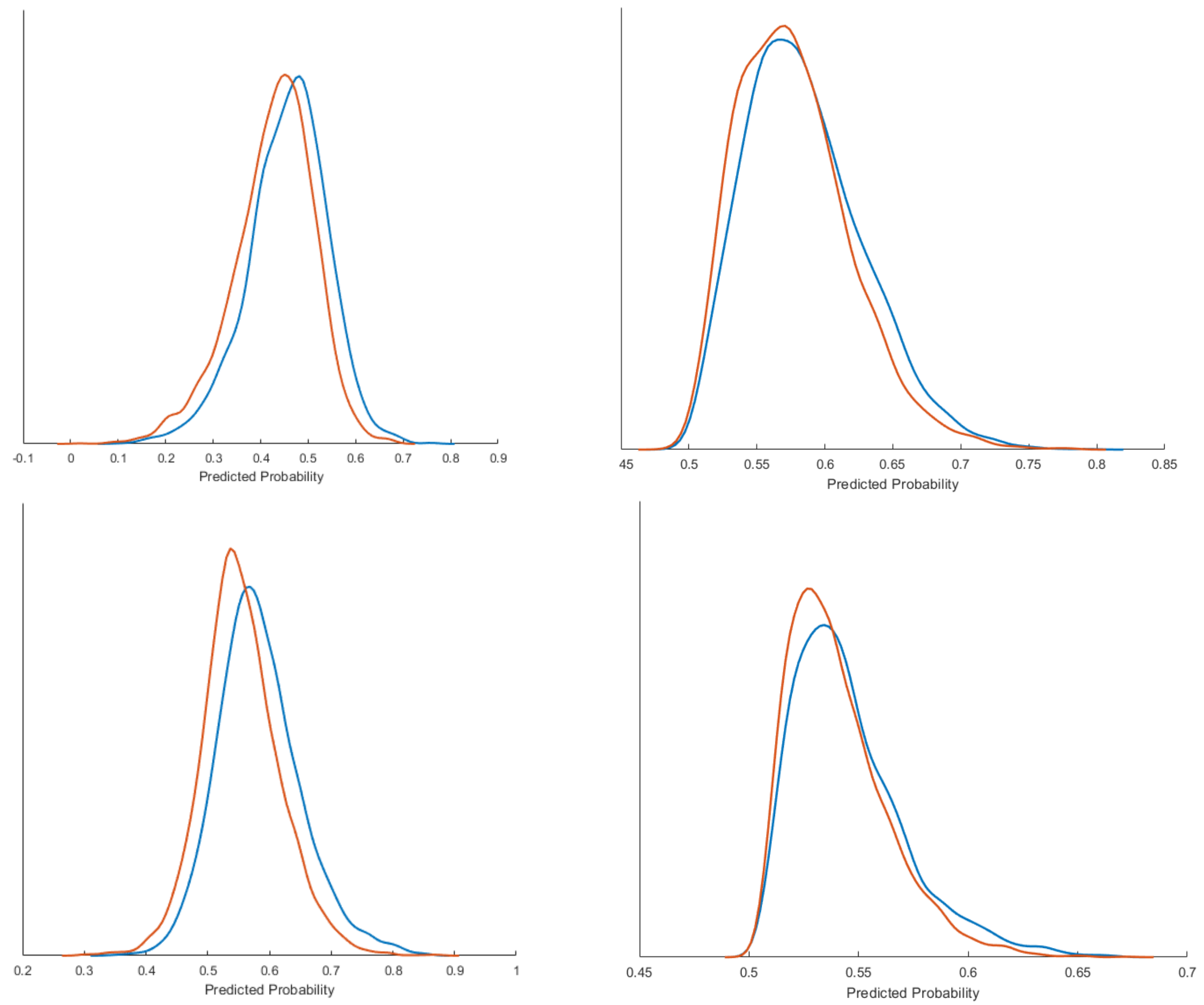


*Note.* The red line shows the distribution of successful stop trials, and the blue shows the distribution of unsuccessful stop trials. Probabilities > 0.5 are classified as unsuccessful, < 0.5 as successful. Single-trial probabilities are calculated by element-wise multiplication between a single-trial time-frequency matrix and the coefficients matrix corresponding to the same component. Next the sum across all elements is calculated. Finally, the sum is converted to log-odds.

**Figure S3**

*Permutation importance for the latency-corrected dataset*

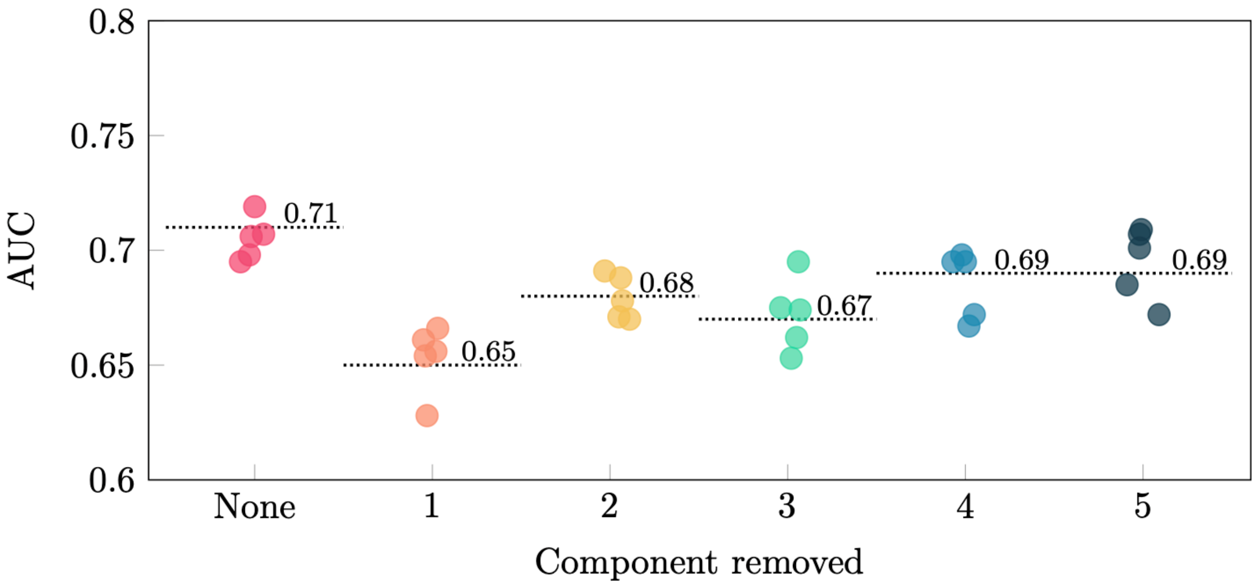


| | None | 1 | 2 | 3 | 4 | 5 |
|---|---|---|---|---|---|---|
| None | | | | | | |
| 1 | * | | * | | * | * |
| 2 | * | | | | | |
| 3 | * | | | | | * |
| 4 | * | | | | | |
| 5 | * | | | | | |

**Supplementary Table 1:** Same as Table 1, but for the latency-corrected dataset

### Minimizing lag between single trials by means of template matching

For each channel:

1. Zero pad data to double the number of samples on each side of the single trial and concatenate all trials.

2. Bandpass filter between 1-10Hz.

3. Reshape back into time x trials and trim padding on either side of the single trial.

4. Construct single trial templates by subtracting the mean from each single trial. Retain samples between 0-500ms where 0 is the stimulus onset.

5. Hann taper each single trial, and z-score normalize. Construct average template by estimating the lag between all single trials. This is done by estimating the normalized autocorrelation and retaining the lag.

6. For each trial, identify the top 5 percentage of trials with the smallest lag. Iterate over the identified trials and circularly shift the identified trials with the estimated lag. For each single trial, calculate the average over the shifted and the current trial.

7. Compute a principal component analysis across the time x trial matrix from step 6. Shrinkage regularization is applied to the covariance matrix with a shrinkage parameter set to 0.1.

8. Retain the top five components, normalize eigenvectors to unit length and calculate component projections.

9. Flip eigenvectors according to the average across single trials estimated in step 5. This involves permuting across the various sign combinations of the eigenvectors and minimizing the Euclidian distance between the z-scored average across the projections against the average across single trials estimated in step 5.

10. The average across the components extracted in step 9 serves as a new template to compare single trials in step 5 against. For each single trial calculate the normalized autocorrelation to the new average and retain the lags.

11. Circularly shift the original time x trial matrix according to the estimated lags in step 10

#### Identifying channels comprising excessive noise involves three steps

1. For each channel, we estimated power-spectral densities using the Matlab function “pwelch”, with 2 second windows, and 50% overlap. Frequencies between 12-40Hz were retained. The average across channels serve as a reference for which each channel’s power spectral density is subtracted. The differences across frequencies are z-scored and the average across these differences are estimated per channel. Channels containing z-scored differences exceeding 1.5 times the interquartile range are flagged as outliers.

2. Calculate the variance for each channel as the diagonal of the covariance matrix estimated from the continuous channel x time data. Define threshold to detect channels whose variance exceeds 1.5 times the interquartile range plus the median of channel variances.
3. The two outlier vectors comprising the flagged channels from both steps were concatenated into a matrix comprising two columns with as many rows as there were channels. The percentage was calculated such that if one channel was flagged as an outlier in both columns, this would result in 100%. Channels flagged by both steps were considered outlier channels.

MATLAB script to detect and attenuate local artifacts following ICA can be shared upon request.
